# Targeting histone deacetylase activity to arrest cell growth and promote neural differentiation in Ewing sarcoma

**DOI:** 10.1101/191700

**Authors:** Bárbara Kunzler Souza, Patrícia Luciana da Costa Lopez, Pâmela Rossi Menegotto, Igor Araujo Vieira, Nathalia Kersting, Ana Lúcia Abujamra, André T. Brunetto, Algemir L. Brunetto, Lauro Gregianin, Caroline Brunetto de Farias, Carol J. Thiele, Rafael Roesler

## Abstract

There is an urgent need for advances in the treatment of Ewing sarcoma (EWS), an aggressive childhood tumor with possible neuroectodermal origin. Inhibition of histone deacetylases (HDAC) can revert aberrant epigenetic states and reduce growth in different experimental cancer types. Here, we investigated whether the potent HDAC inhibitor, sodium butyrate (NaB) has the ability to reprogram EWS cells towards a more differentiated state and affect their growth and survival. Exposure of two EWS cell lines to NaB resulted in rapid and potent inhibition of HDAC activity (1 h, IC_50_ 1.5 mM) and a significant arrest of cell cycle progression (72 h, IC_50_ 0.68-0.76 mM), marked by G0/G1 accumulation. Delayed cell proliferation and reduced colony formation ability were observed in EWS cells after long-term culture. NaB-induced effects included suppression of cell proliferation accompanied by reduced transcriptional expression of the *EWS-FLI1* fusion oncogene, decreased expression of key survival and pluripotency-associated genes, and re-expression of the differentiation neuronal marker ²III-tubulin. Finally, NaB reduced c-MYC levels and impaired survival in putative EWS cancer stem cells. Our findings support the use of HDAC inhibition as a strategy to impair cell growth and survival and to reprogram EWS tumors towards differentiation. These results are consistent with our previous studies indicating that HDis can inhibit the growth and modulate differentiation of cells from other types of childhood pediatric tumors possibly originating from neural stem cells.

## Introduction

Ewing sarcoma (EWS), a highly aggressive bone and soft tissue cancer, is the second most common primary solid bone malignancy in children and young adults [1]. Despite advances in multimodal therapy, patients with the disease have a poor prognosis, with a survival rate of 50 – 65% at 5 years and less than 30% for metastatic or refractory tumors [2]. EWS tumors typically harbor a specific genetic alteration characterized by a chromosomal translocation resulting in fusions between the EWS RNA Binding Protein 1 (*EWSR1*) gene and one of the several *ETS* family genes (most frequently *FLI-1*) which is present in 85% of cases [1,3]. EWS tumors are poorly differentiated and their cell of origin remains elusive. Evidence indicates that EWS shares genetic features and may arise from developing neural crest cells, and has potential for neural differentiation [4-7]. It is also possible that EWS derives from mesenchymal stem cells [8-11]. Experiments investigating clear cell sarcoma, a cancer type defined by *EWS-ATF1*, the fusion product of a balanced chromosomal translocation between chromosomes 22 and 12, have shown that the differentiation state of cells of origin impacts tumor characteristics [12].

EWS is a relatively genetically stable, fusion oncogene-driven cancer, harboring few somatic mutations [13]. However, as with other solid tumors of childhood, many epigenetic alterations are likely crucial for EWS tumorigenesis [14]. EWSR1-FLI-1 acts as an aberrant transcription factor that induces epigenomic reprogramming involving changes in chromatin remodeling through acetylation and methylation, resulting in repression of tumor suppressors and oncogene activation [15, 16]. The chromatin state in EWS is strikingly similar to that found in bone-marrow-derived mesenchymal stem cells. Increased chromatin accessibility in stem cells may lead to a state that facilitates oncogenic alterations induced by EWSR1-FLI-1, suggesting a stem cell origin for EWS [17].

A balance between the opposing activities of histone acetyltransferases (HATs) and deacetylases (HDACs) is key in regulating gene expression. Histone acetyltransferases (HATs) control histone acetylation activity through the transfer of acetyl groups to the amino-terminal lysine residues of histones, thus increasing transcriptional activity. In contrast, histone deacetylases (HDACs) remove acetyl groups, favoring chromatin condensation and repressing gene expression [18]. HDAC inhibitors (HDi) represent a class of experimental antineoplastic agents that target aberrant epigenetic alterations found in cancer [19, 20]. Sodium butyrate (NaB) is an HDi widely used as an experimental tool in cellular studies. It is short fatty acid that acts as a potent class I and IIa HDAC inhibitor inducing arrest in proliferation of cancer cells by increasing histone acetylation. In mammalian cells, NaB promotes hyperacetylation of the histones H3 and H4, resulting in chromatin decondensation and increased transcriptional activity [21]. In EWS cells, we have previously shown a synergistic antitumor effect of NaB combined with commonly chemotherapeutic drugs [22]. Moreover, the HDi suberoylanilide hydroxamic acid (SAHA) showed a synergistically enhanced antitumor activity upon combination with etoposide [23], and HDis NaB, SAHA or entinostat or MS-275 as monotherapies or combined with TRAIL display an additive cytotoxic effect in EWS cells [24].

Here, we show that inhibition of global HDAC activity by NaB inhibits multiple pathways involved in cell proliferation, survival and pluripotency. In addition, NaB exposure stimulates some morphologic and biochemical changes that might be consistent with neural differentiation in EWS cells.

## Materials and Methods

### Cell Lines and Cell Culture

Human EWS cell lines SK-ES-1 and RD-ES-1 obtained from American Type Culture Collection (Rockville, MD, USA) were grown in RPMI-1640 medium (Gibco-BTL, Carlsbad, CA, USA), containing 0.1% fungizone^®^ (250 mg/kg; Invitrogen Life Technologies, São Paulo, Brazil), 4 mg/ml gentamicin and 10% fetal bovine serum (FBS, Gibco^®^ by Thermo Fisher Scientific, Life Technologies, Brazil). Cells were cultured at 37 °C in a humidified incubator under 5% CO_2_.

### Drug Treatment

NaB (Sigma Aldrich - St. Louis, MO, USA) was diluted in sterile ultrapure water in a stock solution of 100 mM. EWS cells (2×10^4^) were plated in 24-well plate and exposed to NaB in concentrations ranging from 0.5 to 5 μM for 72 h. For calculation of IC_50_, data were fitted in a dose response curve (Graphpad Prism v. 6.0) using the equation: *y = min + (max – min)/(1 + 10^((LogIC50 – x)*Hillslope + Log ((max – min)/(50-min) − 1)))*.

### Cell Viability

Cells were exposed to NaB as described above. Both cells in the supernatant and adhered cells were detached, collected, centrifuged and washed with PBS twice. To assess viability, cells were incubated with 1 μg/ml propidium iodide (PI) (Sigma Aldrich) in PBS at 4 °C in the dark. PI uptake was assessed by flow cytometry analysis using an Attune Acoustic focusing cytometer (Applied Biosystems, Thermo Fisher Scientific, USA). Data was analyzed using Attune Cytometric Software version 1.2.5. Four individual replicates were performed.

### Cell Cycle

To assess cell cycle, the EWS cells were cultured in 24-well plates under the same conditions of described above, and then detached, centrifuged and washed with PBS twice. After, the cells were ressuspended in 50 µg/ml propidium iodide (Sigma-Aldrich, St. Louis, Mo., USA) in 0.1% Triton X-100 in 0.1% sodium citrate solution containing 5 µg/ml RNase A. One million cells were stained in 1.0 ml of the PI cell cycle solution for 5 min. followed by assessment on an Attune Acoustic focusing cytometer by Applied Biosystems (Thermo Fisher Scientific, USA) analysis. In each sample 20,000 cells were analysed. Data was analyzed using FlowJo Cytometric Software version 10.1. Three individual experiments were performed.

### Cumulative Population Doubling

Cells were plated [2×10^4^ cell/mL] and exposed to NaB, in the same conditions described above, in quadruplicates. Every 4 days, supernatant and adhered cells were collected, quantified by PI uptake and re-plated. The Population Doubling (PD) level was calculated to determine the proliferation potential of the cell lines SK-ES 1 and RD-ES unexposed versus NaB exposed cells. The PD at each passage was calculated using the equation PD = (log Xe – log Xb)/log2 and doubling time was calculated using DT = T*ln2/ln(Xe/Xb). In both cases ‘Xe’ is the cell number at the end of the incubation time, ‘Xb’ is the cell number at the beginning of the incubation time and T is the number of days between time points, as described in the ATCC^®^ Animal Cell Culture Guide. Data is displayed as cumulative PD (CPD), calculated as the sum of all previous PDLs.

All the CPD parameters were analyzed according to Silva and colleagues [25]. Briefly, Relative end CPD (RendCPD) quantifies the end point of cell proliferation analysis, and is obtained by the ratio between the final CPD of cells exposed to drug and control cells. To determine the global effect after the treatment intervention, the relative Area Under Curve (rAUC) was calculated (Graphpad Prism v. 6.0 software) using the lower CPD of the initial population to obtain the minimal threshold. RPR (Relative Proliferation Rate) and RTCT (Relative Time to Cross a Threshold), linear regression for both exposed and control groups was calculated using the higher angular coefficients of the last two or more CPD intervals. RPR is the ratio of the angular coefficients of exposed and control groups. RTCT was determined by the ratio of between the control and exposed cells Time to Cross the Threshold (TCT), *i.e.*, a threshold set on a CPD value where both treatment and control conditions present the highest velocity of proliferation. Long-term survival index (LSI) is calculated by the average of the four parameters described above.

### Colony Formation

Cells were exposed to NaB for 72h and 1 × 10^2^ cells per well were re-plated in 6-well plates and cultured for 10 days with the RPMI medium being changed every 3 days. Cells were then fixed with methanol, followed by staining with 0.5% crystal violet. The stained colonies images were scanned and analyzed using the cell counter plugin on Image J software (NIH, Bethesda, MD, USA).

### Tumorsphere Culture

Cells were dissociated with trypsin-EDTA into single cell suspension and seeded at 2 × 10^3^ cells/well in 6-well plates. The cells were cultured in a serum-free tumorsphere (TS)-inducing medium, containing DMEM-F12 (1:1) supplemented with 2% B27 supplement (Gibco, Invitrogen, CA, USA), 20 ng/mL recombinant human EGF (Sigma-Aldrich, MO, USA), 20 ng/mL human leukemia inhibitor factor (Thermo Fisher Scientific, Life Technologies), 10 IU/mL (5 mg/mL) heparin (Roche, Mannheim, Germany) and antibiotics as described by Leuchte and colleagues [26]. The media was changed every 4 days. After 8 – 9 days, tumorspheres were dissociated with a non-enzymatic solution containing 1 mM EDTA, 40 mM Tris-HCl and 150 mM NaCl, and re-plated in a 96-well plate to evaluate their capacity to self-renew through secondary EWS tumorsphere formation. Tumorspheres with at least 80μm were analyzed and quantified by inverted phase microscopy (Leica Microsystems, Mannheim Germany). The effect of NaB on tumorsphere formation ability across 7 days was examined.

### HDAC Activity

Cells (1 × 10^4^) were seeded in a 96-black well plate/clear flat-bottom (Greiner-bioone, Frickenhausen, DE) and exposed to NaB, at the same concentrations described above, for 1 or 3 h. HDAC enzymatic activity was measured by *In Situ* HDAC Activity Fluorometric Assay kit (EPI003, Sigma-Aldrich, St. Louis, MO, USA) according to the manufacturer’s instructions. Three individual experiments were performed. HDAC enzyme activity was calculated across the groups as pmol/min and the percent of HDAC activity was corrected by control (unexposed cells).

### Immunofluorescence

For immunofluorescent staining, cells were plated on the glass coverslips, pre-coated with 2 μg/ml of fibronectin. After 72 h of exposure to NaB, cells were fixed in 4% PFA for 15 minutes at room temperature. After two additional washes, the fixed cells were incubated with blocking buffer (1X PBS containing 3% BSA and 0.3% Triton X-100) for 1 h at room temperature, followed by an overnight incubation at 4°C with the following primary antibodies: Histone 3 phos-S10 (1:200, Cell Signaling), Trkb (1:100, Santa Cruz). After washing with PBS twice, cells were incubated with secondary antibodies conjugated with either Alexa fluor-488 (1:800; Invitrogen) or Rhodamine (1:300, Invitrogen) at room temperature for two h. After three additional washes, DNA was stained with 4 μg/ml of Hoechst 33342 (Invitrogen) for 15 minutes at room temperature. After washing, slides were mounted with Prolong Gold anti-fade reagent. Slides were viewed with Leica TCS SP5 Confocal Laser Scanning Microscope, using Leica LAS AF Lite 2.0.2 software to acquire images. For quantitative analysis, was calculated by the percent of mean intensity corrected by the control (unexposed cells). For each group, at least five random images were taken, and the results were determined from at least three independent experiments.

### Real Time qPCR

Total RNA was extracted from EWS cells using TRIzol (Invitrogen, Carlsbad, CA, USA). Prior to RT-PCR, the samples were treated with DNase I (Promega Corporation, WI, USA). Total RNA was quantified with NanoDrop (Thermo Fisher Scientific, DE, USA). SuperScript^TM^ First-Strand Synthesis System for RT-PCR (Invitrogen, Carlsbad, CA, USA) enabled first strand synthesis. The mRNA expression levels of target genes (*EWS-FLI1*, TrkB, OCT3/4) were quantified using KiCqStart^TM^ SYBR Green qPCR ReadyMix^TM^, with ROX^TM^ (Sigma-Aldrich, St. Louis, MO, USA) and calculated using the ΔCT method from triplicate reactions, with the levels of gene normalized to the relative Ct value of Gapdh. Cycling parameters were as follows: 95 °C for 2 mins, followed by 40 cycles of denaturation at 95 °C for 3 secs, annealing at 60 °C for 15 secs, and extension at 72 °C for 30 secs. The primer sequences used are shown in Supplementary Table S1 online.

### Western Blot

Proteins were separated using 8 – 14% SDS Tris-glycine gels and transferred onto a polyvinylidene difluoride membranes. Membranes were blocked with 5% fat-free milk and incubated with antibodies against BDNF (H117, SC-20981, Santa Cruz, CA, USA), NGF (H-20, sc-548, Santa Cruz, CA, USA), pERK (E-4 sc-7383, Santa Cruz, CA, USA), ERK1 (K-23, sc-94, Santa Cruz, CA, USA), pAKT (S473) (SAB4300042, Sigma Aldrich, MO, USA), AKT (pan) (C67E7, #4691, Cell Signaling Technology, MA, USA), OCT3/4 (sc-5279, Santa Cruz, CA, USA), NANOG (M-149, sc-33760, Santa Cruz, CA, USA), c-MYC (D84C12 – Cell Signaling Technology, MA, USA), KLF4 (H-180, SC-20691, Santa Cruz, CA, USA), ALDH1A1 (ab23375, Abcam, Cambridge, UK), BMI-1 (D20B7, #6964 – Cell Signaling Technology, MA, USA), Cyclin D1 (D71G9 – Cell Signaling Technology, MA, USA), βIII-Tubulin (D71G9 – Cell Signaling Technology, MA, USA) and ACTB (A2228, Sigma Aldrich, MO, USA) used as loading control. Incubation for 1 h with appropriate horsedish peroxidase-conjugated secondary antibody (Santa Cruz) at RT was performed. Chemiluminescence was detected using ECL Western Blotting substrate (EMD Millipore, DE) and analyzed by ImageQuant LAS500 (GE Healthcare Life Sciences, UK). Densitometric analyses were performed using Image J (NIH, MD, USA). Relative densitometric unit (RDU) was calculated by the normalization of interest protein level to the housekeeping genes β-actin (ACTB) or Histone 3 (H3), or ERK1 and AKT for pERK and pAKT (S473), respectively. All treatment conditions were corrected by control groups (non-exposed cells). Three individual replicates were performed.

### Statistical Analysis

For statistical analysis, GraphPad Prism version 6.0 was used. Each assay was performed at least three times in biologically independent assays. The significance of differences in mean ± standard error values was analyzed by one-way ANOVA and Tukey’s multiple comparison test or Student’s *t*-test; *p* values of less than 0.05 were considered statistically significant.

## Results

### Inhibition of HDAC Activity Reduces *EWS-FLI1* Fusion Oncogene Transcriptional Expression and Induces Growth Arrest in EWS Cells

To examine the effect of NaB on HDAC activity in EWS cells, we exposed SK-ES1 and RD-ES cells to NaB for 1 and 3 h. HDAC activity was significantly reduced after 1 h when NaB doses higher than 1 mM were used, resulting in a target IC_50_ of approximately 1.5 mM (Fig. 1a, 1d). HDAC activity in cells exposed to NaB for 3 h was not as effectively reduced in comparison to 1 h treatments, and a significant decrease was observed only in RD-ES cells exposed to 2 mM of NaB (Supplementary Fig. S1), suggesting that NaB optimally inhibits HDAC activity in EWS cells at a kinetic rate of 1 h.

**Fig. 1.**
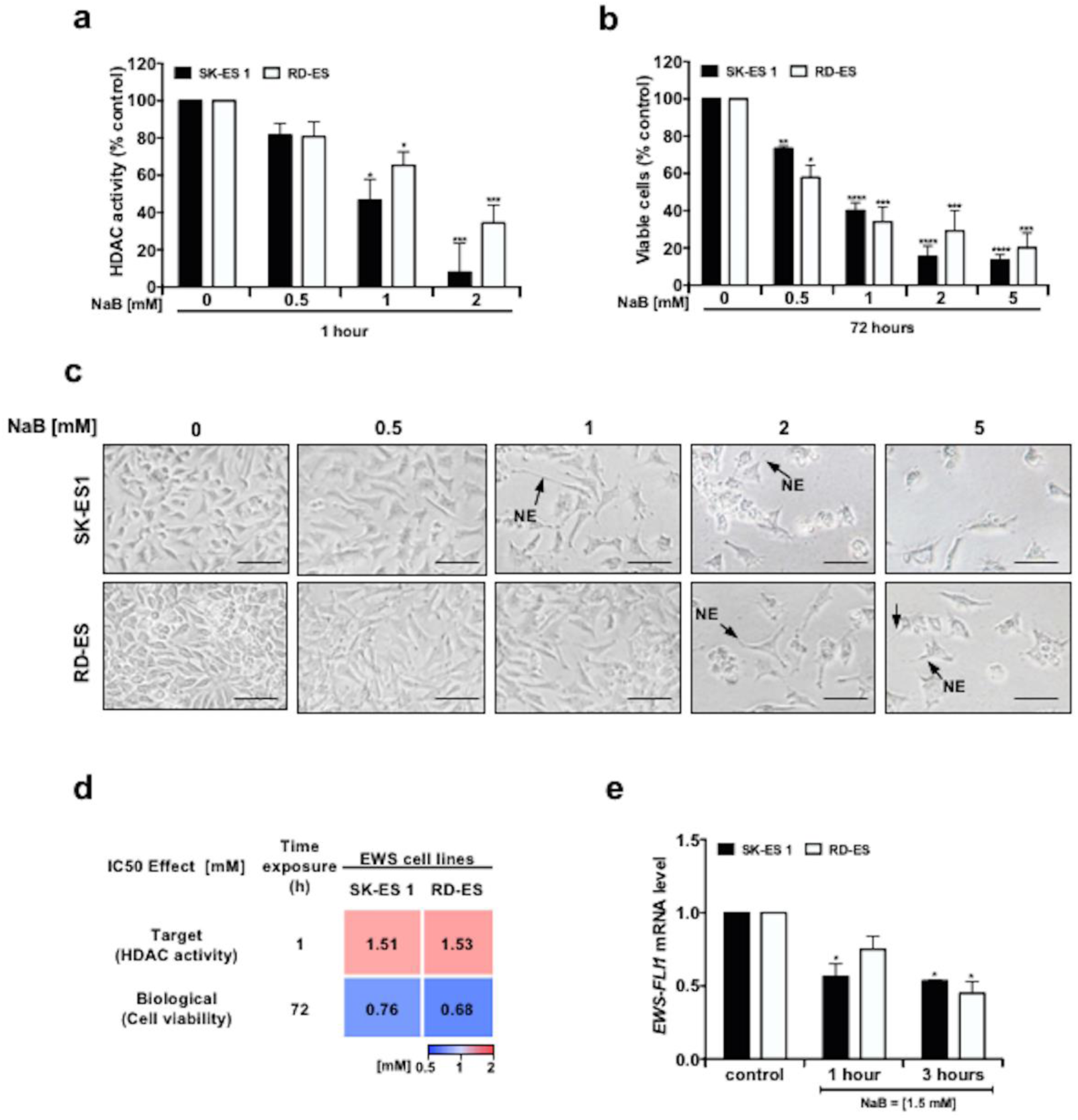
HDAC Inhibition by NaB hinders the growth of EWS cells. **a** HDAC activity (%) in EWS cell lines SK-ES 1 and RD-ES after 1 h of exposure to NaB; *N = 4* independent experiments. **b** Percent of viable SK-ES 1 and RD-ES cells after 72 h of exposure to NaB; *N = 4* independent experiments. **c** Morphology of EWS SK-ES 1 (upper panel) and RD-ES (lower panel) cells after 72 h of NaB exposure; black arrows indicates neurite-like extensions (NE). Scale bar: 50 μm. **d** Heat map showing the target IC_50_ calculated by the percentage of HDAC activity in cells exposed to NaB for 1 h, and biological IC_50_ calculated by the percent of viable cells exposed to NaB for 72 h**. e** *EWS-FLI1* mRNA levels in EWS cell lines SK-ES 1 and RD-ES after 1.5 mM of NaB exposure for 1 and 3 h. Gene expression levels were normalized by gapdh mRNA level and corrected by untreated EWS cells; *N = 3* independent experiments. Data in the graphs are shown as mean ± SEM; *\* p <* 0.05, *\*\* p <* 0.01, *\*\*\* p <* 0.001, *\*\*\*\* p <* 0.0001 *vs*. controls.

In order to evaluate the biological effects of HDAC inhibition, we exposed EWS cells to varying concentrations of NaB (0.5 – 5 mM) for 72 h. Cell viability was potently reduced in both cell lines (Fig. 1b). Interestingly, cells exposed to NaB showed a change in morphology accompanied by the appearance of short neurite-like extensions (Fig. 1c). At 72 h, the biological IC_50_ of NaB was 0.76 and 0.68 mM for SK-ES 1 and RD-ES EWS cell lines, respectively (Fig. 1d). Moreover, to examine whether HDAC inhibition could alter the expression of the fusion oncogene, we designed primers flanking the break point of *EWS-FLI1* fusion (type 2) and evaluated its transcriptional expression level after NaB exposure in comparison with untreated EWS cells. HDAC inhibition in EWS cells resulted in an approximately 2-fold decrease of *EWS-FLI1* transcriptional expression after treatment with 1.5 mM NaB at 1 and 3 hours for SK-ES 1 cells, and 3 hours for RD-ES cells (Fig. 1e).

Next, we verified whether inhibition of HDAC activity by NaB would change cell cycle distribution. HDAC inhibition resulted in a significant alteration in EWS cell cycle featuring an accumulation of cells in the G0/G1 phase 35 h after NaB exposure. In the SK-ES1 EWS cell line, we also observed a significant decrease in the S and G2/M phases of the cell cycle, whereas in the RD-ES cell line there was a significant reduction in polyploidy (Fig. 2a). To determine whether HDAC inhibition disrupts histone 3 phos-S10, a chromosome condensation marker during mitosis, we immunostained EWS cell lines exposed to NaB for 72 h against anti-H3 phos-S10 plus anti-Alexa488, measured by laser confocal microscopy. As expected, H3 phos-S10 immunolocalized to the nucleus in both EWS cell lines (Fig. 2b). In addition, we observed that both EWS cell lines exposed to higher doses of NaB (2 and 5 mM) showed a reduced level of H3 phos-S10, whereas there was no significant change at lower doses (0.5 and 1 mM) compared to untreated cells (Fig. 2c). Given that Cyclin D1 is a G1-phase regulator protein overexpressed in EWS, we investigated whether HDAC could regulate Cyclin D1 expression after 72 h of NaB exposure. SK-ES 1 cells exposed to 0.5 or 1 mM of NaB showed an approximately 1.5-fold change decrease in Cyclin D1 levels, and 1 mM NaB also decreased Cyclin D1 levels also in RD-ES cells (Fig. 2d). The results suggest that HDAC activity inhibition induces growth arrest in EWS cells.

**Fig. 2.**
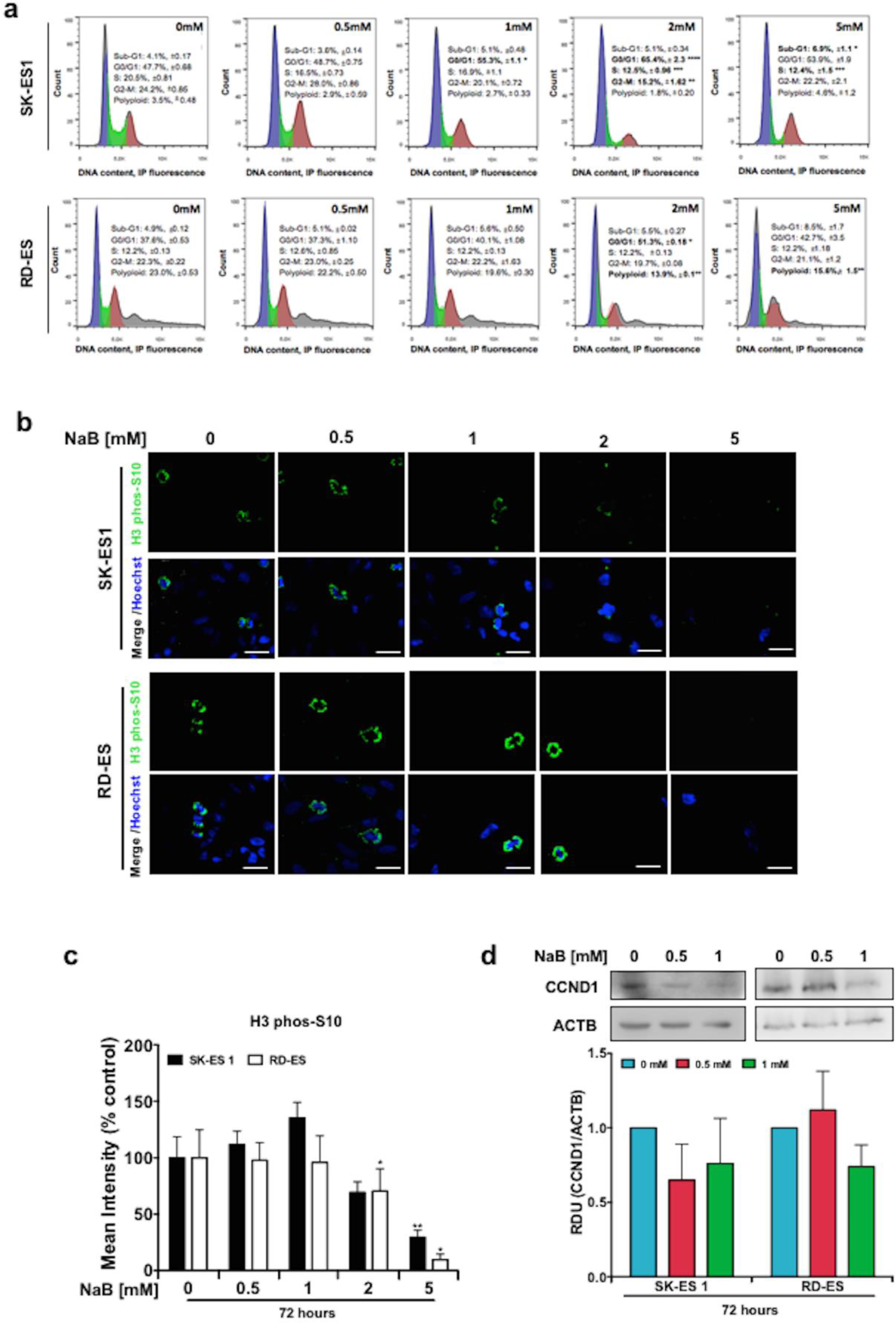
Cell cycle changes induced by HDAC inhibition in EWS cells. **a** Cell cycle distribution of EWS SK-ES 1 (upper panel) and RD-ES (lower panel) cells after 35 h of NaB exposure; *N = 3* independent experiments. **c** Representative images of H3 phos-S10 level (immunofluorescence) from SK-ES 1 (upper panel) and RD-ES (lower panel) cells exposed to NaB for 72 h. **c** Percent of H3 phos-S10 mean intensity (immunofluorescence) from EWS cell lines exposed to NaB for 72 h; *N* = 3 independent experiments. **d** Western blot (upper panel) of Cyclin D1 protein levels in EWS cells exposed to NaB for 72 h, and Relative Densitometric Unit (RDU) analysis (lower panel) normalized by ACTB, and corrected by control. Data in the graphs are shown as mean ± SEM; *\* p < 0.05, ** p < 0.01, *** p < 0.001, **** p <* 0.0001 *vs*. controls.

### HDAC Inhibition Abrogates Cell Proliferation and Survival Pathways in EWS Cells

To investigate molecular mechanisms associated with the NaB-induced decrease in EWS cell growth, we examined the expression of key proteins that control cell survival and proliferation. After exposure to either 0.5 or 1.0 mM NaB, a decrease in the levels of survival-promoting neurotrophins, brain-derived neurotrophic factor (BDNF) and nerve growth factor (NGF) were observed (Fig. 3a). We also immunostained tropomyosin receptor kinase B (TrkB) in EWS cell lines against anti-TrkB, plus anti-Rhodamine. As expected, TrkB immunoreactivity was detected in the cytoplasm. In addition, we observed that TrkB levels were reduced in a dose-dependent manner in both cell lines after NaB exposure (Fig. 3b, 3c). Consistently with this finding, we also observed a decrease in protein levels of downstream targets, phosphorylated extracellular-regulated kinase (pERK), but not total ERK1 levels, pAKTS473, and total AKT. A significant reduction of AKT was observed only in RD-ES cells treated with 1.0 mM NaB when pAKTS473 over total AKT levels were considered (Fig. 3d). Essentially similar effects were observed in RD-ES cells, however only at the 1.0 mM dose of NaB (Fig. 3d). Analysis of relative densitometric units related to the immunoblots is depicted in Fig. 3e. These results support the possibility that signaling mediated by neurotrophins, ERK and AKT are involved in mediating the inhibitory effects of NaB in EWS cells.

**Fig. 3.**
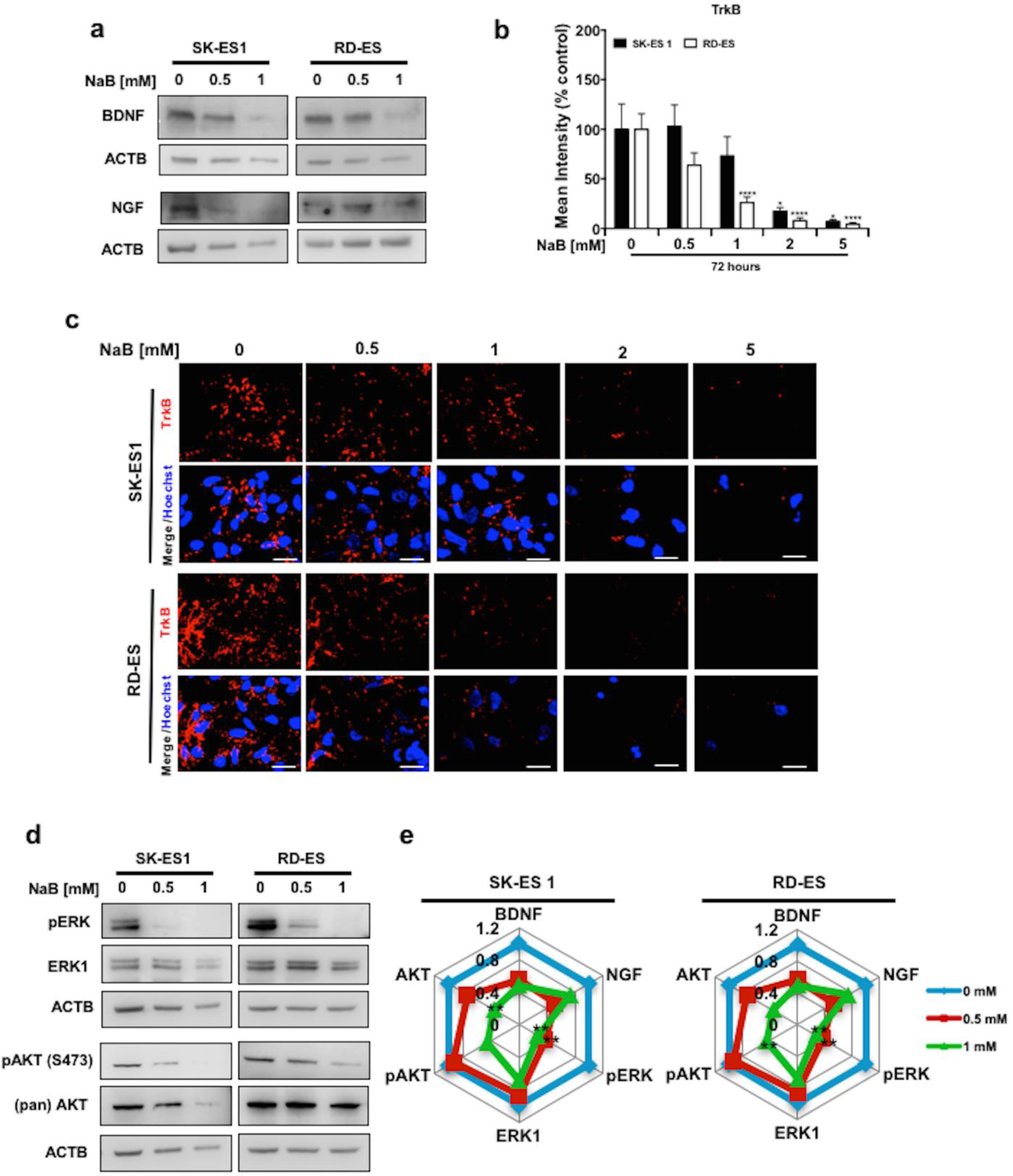
HDAC activity inhibition decreases components of signaling pathways related to proliferation and survival in EWS cells. **a** Western Blot of BDNF and NGF from SK-ES 1 (left) and RD-ES (right) cells exposed to NaB for 72 h. **b** Percent of TrkB mean intensity (immunofluorescence). **c** Representative images of TrkB level (immunofluorescence) from SK-ES 1 (upper panel) and RD-ES (lower panel) cells exposed to NaB for 72 h. **d** Western Blot analysis of pERK, ERK1, pAKT(Ser473), AKT in SK-ES 1 (left panel) and RD-ES (right panel) cells. **e** Radar graph showing Relative Densitometric Unit (RDU) analysis were normalized by ACTB, except for pERK and pAKT(Ser473), which were normalized by total ERK1 and AKT, respectively, and corrected relative to control cells. Data in the graph are shown as mean ± SEM from three independent experiments; *\* p <* 0.05, *\*\* p <* 0.01, *\*\*\*\* p <* 0.0001 *vs*. controls.

### Persistence of the Anti-Proliferative Effect Induced by HDAC Inhibition in EWS Cells

We performed a cumulative population doubling (CPD) of EWS cells to examine the persistence of the anti-proliferative effect of NaB. After 72 h of treatment, cells were maintained for 16 days in drug free media, as shown in Fig. 4a. Significant delays in cell proliferation rate were found with 2 and 5 mM of NaB (Fig. 4b). Because the number of cells was too small, we modeled the growth from day 19 to day 35 using a mathematical prediction as described by Silva et al. [25]. We were able to predict whether EWS cells exposed to the higher doses of NaB over 35 days would have a persistent reduction in the ability to proliferate. Lower doses (0.5 and 1 mM) of NaB resulted in a gain in cell proliferation by day 20 which by 35 days was even higher than our prediction (Fig. 4b). It is possible that only a resistant subset of EWS cells could restore cell proliferative ability to an equal or higher degree compared to controls.

**Fig. 4.**
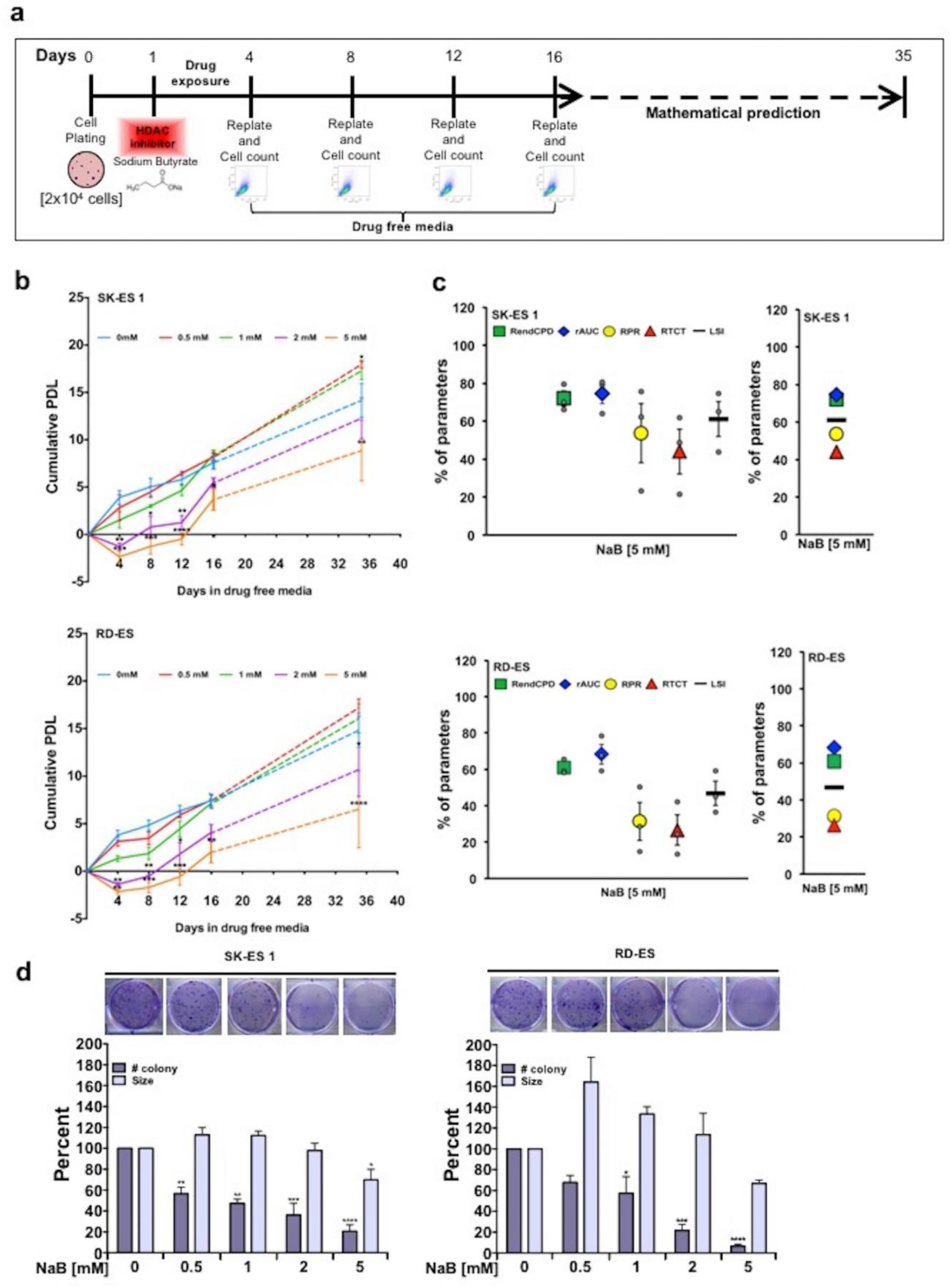
Persistent inhibition of cell proliferation rate and colony formation ability in EWS cells treated with NaB. **a** Cumulative Population Doubling (CPD) *in vitro* assay schematic and timeline. **b** Cumulative CPD from SK-ES 1 (upper panel) and RD-ES (lower panel) EWS cells after 72 h of NaB treatment plus 16 days in a drug free media, plus 19 days of a mathematical prediction (total 35 days); *N = 3* independent experiments. **c** CPD profile from SK-ES 1 (upper) and RD-ES (lower) EWS cells after 72 h of NaB (5 mM). *(Green square – RendCPD, Blue diamond rAUC, yellow circle – RPR, red triangle – RTCT, black line - LSI)*. **d** Colony formation in SK-ES1 (left panel) and RD-ES (right panel) cells after 72h of NaB exposure plus 10 days in a drug free media, measured by percent of colony number and size; *N = 3* independent experiments. Data in the graphs are shown as mean ± S.E.M; *\* p <* 0.05, *\*\* p <* 0.01, *\*\*\* p <* 0.001, *\*\*\*\* p <* 0.0001 *vs*. controls.

We then went on to perform a CPD profile analysis that examines long-term changes in cell population size. On the basis of CPD graphs from cell counting we calculated four different parameters: relative end CPD (RendCPD) to quantitatively assess the end point of cell proliferation analysis; relative area under curve (rAUC) to determine a global effect after treatment intervention; relative time to cross a threshold (RTCT) to measure the delay of the cell population that recovers growth after treatment; and relative proliferation rate (RPR) to quantify the relative regrowth velocity of the cells which survived after treatment intervention. The average of all parameters described above is termed the long-term survival index (LSI). HDAC activity inhibition in both cell lines, particularly after treatment with 5 mM NaB, resulted in a robust decrease in RPR and RTCT parameters (Fig. 4c), except the RPR parameter in which there was no significant change in SK-ES 1 cells (data not shown). These findings indicate that the effects of NaB can persist in the long-term to delay the regrowth of the EWS cell population.

Finally, we performed a colony formation assay for 10 days in drug free media, after a 72-h exposure to NaB. There was a significant reduction in the number of colonies in both cell lines after exposure to NaB. Moreover, there was a significant decrease in the size of the colonies in SK-ES 1 cells after exposure to 5 mM NaB (Fig. 4d). These results indicate that the decrease in cell growth is also associated with the cells’ colony-forming capability.

### HDAC inhibition reduces the expression of pluripotency-associated genes

Previous studies have shown that the EWSR1/FLI-1 oncoprotein stimulates the pluripotency gene BMI-1, thus influencing EWS cell self-renewal [7]. We sought to evaluate by immunoblot analysis whether the expression of key pluripotency transcriptional regulators NANOG, c-MYC, OCT3/4, KLF4, BMI-1 and ALDH1A1 were altered because of HDAC inhibition by NaB. Both SK-ES 1 and RD-ES cells showed a 2–3-fold reduction in the levels of pluripotency-associated genes OCT3/4, c-MYC and BMI-1 compared to untreated control cells. The levels of KLF4, NANOG and ALDH1A1 decreased in SK-ES 1 cells but were relatively unchanged or increased in RD-ES cells (Fig. 5a, 5b; Supplementary Fig. S2, S3). Thus, SK-ES 1 cells show higher sensitivity to NaB regarding inhibition of pluripotency-associated genes.

**Fig. 5.**
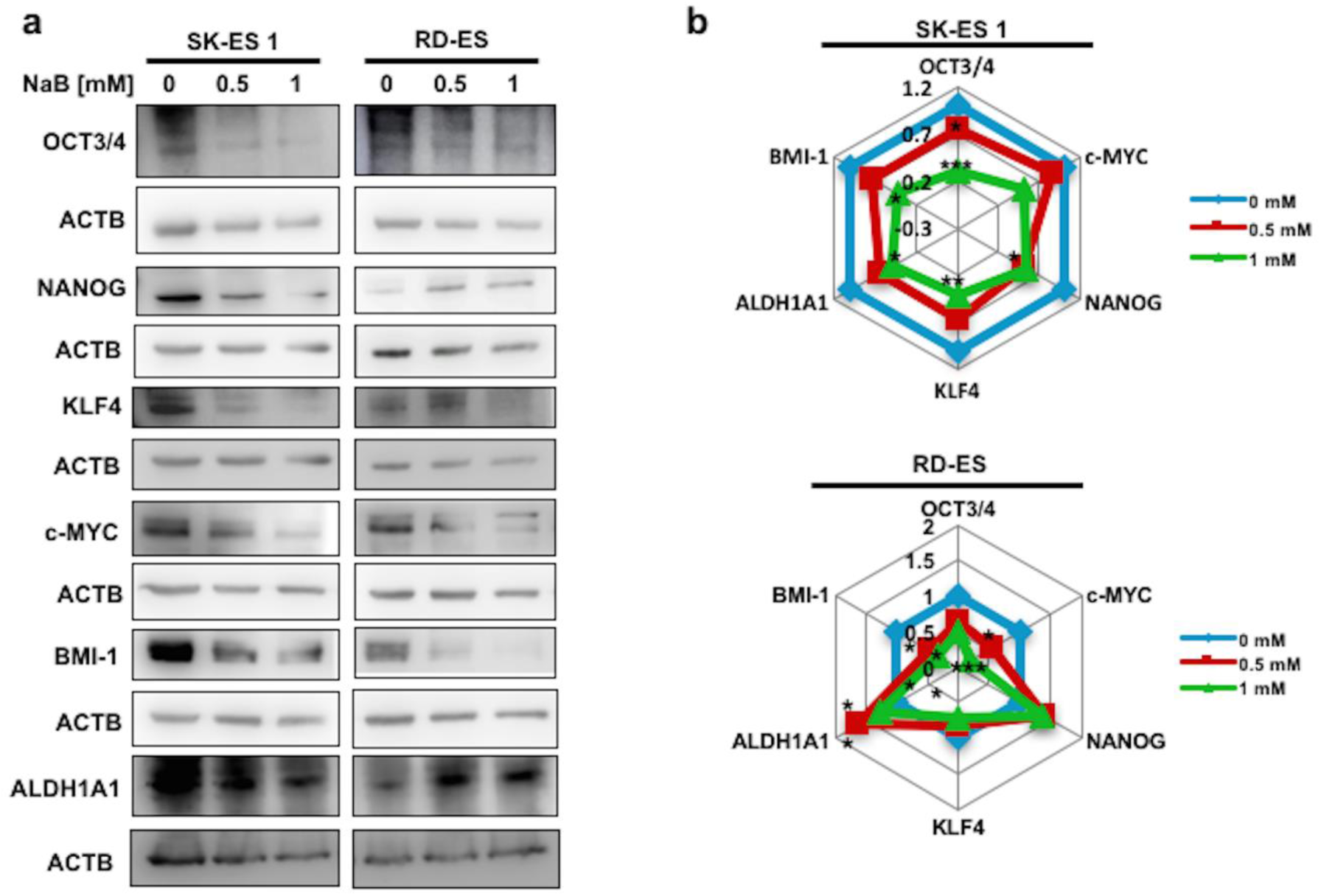
Changes in the expression of pluripotency-associated genes induced by HDAC inhibition in EWS cells. **a** Western Blot analysis of pluripotency-associated genes in SK-ES 1 (left panel) and RD-ES (right panel) cells after 72 h of exposure to NaB. **b** Radar graph showing Relative Densitometric Unit (RDU) analyses normalized by ACTB level, and corrected by control; *\* p < 0.05, ** p < 0.01, *** p < 0.001 vs*. controls.

Given that we observed morphological changes in EWS cells after NaB exposure, we asked whether the SSC-A parameter that measure complexity and granularity in flow cytometry is altered in EWS cell population after the pharmacological intervention by NaB. Interestingly, both EWS cell lines exposed to NaB showed an approximately 2-fold increase in complexity or granularity measured by the SSC-A mean parameter in flow cytometry (Fig. 6a, 6b). These data provide a measurement parameter related to cell morphology alterations along EWS cell population. In addition, this alteration was accompanied by cell morphology changes with formation of neurite-like extensions (Fig. 6c). Furthermore, we assessed the expression level of the neural differentiation marker β-III Tubulin (TUBB3) (Fig. 6d). We observed an increase in TUBB3 in cells exposed to HDAC activity inhibition by NaB, with a 2-fold enhancement in protein levels compared to controls. These findings suggest that HDAC activity inhibition may act to reduce the expression of a set of genes related to pluripotency in EWS cells.

**Fig. 6.**
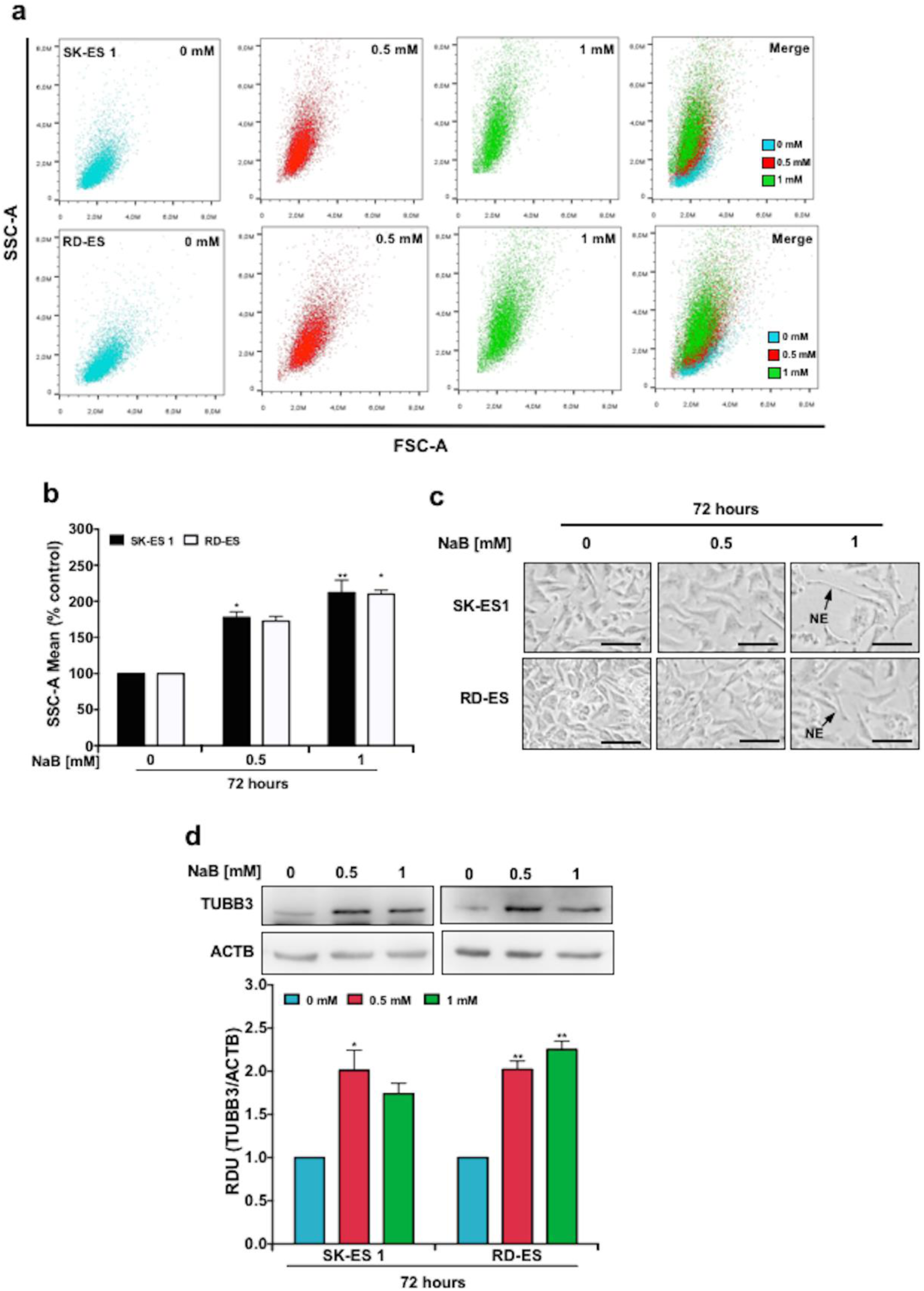
Cell complexity and differentiation of EWS cells after HDAC inhibition. **a** Representative scatter plot from SK-ES 1 (upper) and RD-ES (lower) exposed to NaB for 72 h, measured by cytometric analysis. **b** Percent of SSC-A mean from EWS cells exposed with the indicated concentrations of NaB for 72 h; *N* = 4 independent experiments. **c** Morphology of SK-ES 1 (upper panel) and RD-ES cells (lower panel) after 72 h of NaB exposure; black arrows indicate neurite-like extensions (NE). Scale bar: 50μm. **d** Western Blot analysis of the differentiation neural marker TUBB3 (upper panel); *N = 3* independent experiments, and Relative Densitometric Unit (RDU) analyses (lower panel) normalized by ACTB, and corrected by control. Data in the graph are shown as mean ± SEM; *\* p <* 0.05, *\*\* p <* 0.01 *vs*. controls.

### HDAC Inhibition Impairs EWS Tumorsphere Formation

Cancer stem cells displaying tumor-initiating properties and treatment resistance ability have been identified and characterized in EWS [27]. Tumorsphere (TS)-forming assays have been widely used as models to study cancer stem cell-enriched cultures [28]. To verify whether HDAC inhibition altered TS-forming ability of EWS *in vitro*, we measured TS-formation efficiency (TFE) and size through optical microscopy after exposure of cells to NaB in a serum-free TS-inducing medium for 7 days (Fig. 7a). The contents of the stem cell marker OCT3/4 and TrkB were increased in cells from TS compared to ML (Fig. 7b, 7e). NaB significantly reduced TFE in both cell lines. In addition, NaB reduced the size of EWS tumorspheres (Fig. 7c, 7d; Supplementary Fig. S4). It is noteworthy that the higher dose of NaB (5 mM) fully impaired the cells’ ability to produce tumorspheres. Interestingly, the levels of BMI-1 were also increased in TS compared to cells in the ML, and NaB significantly reduced the levels of c-MYC in TS cells (Fig. 7d, 7e). These findings indicate that OCT3/4, TrkB, and BMI-1 are increased in EWS TS cultures, and suggest that HDis can repress c-MYC expression in EWS cancer stem cells and decrease their ability to expand and survive.

**Fig. 7.**
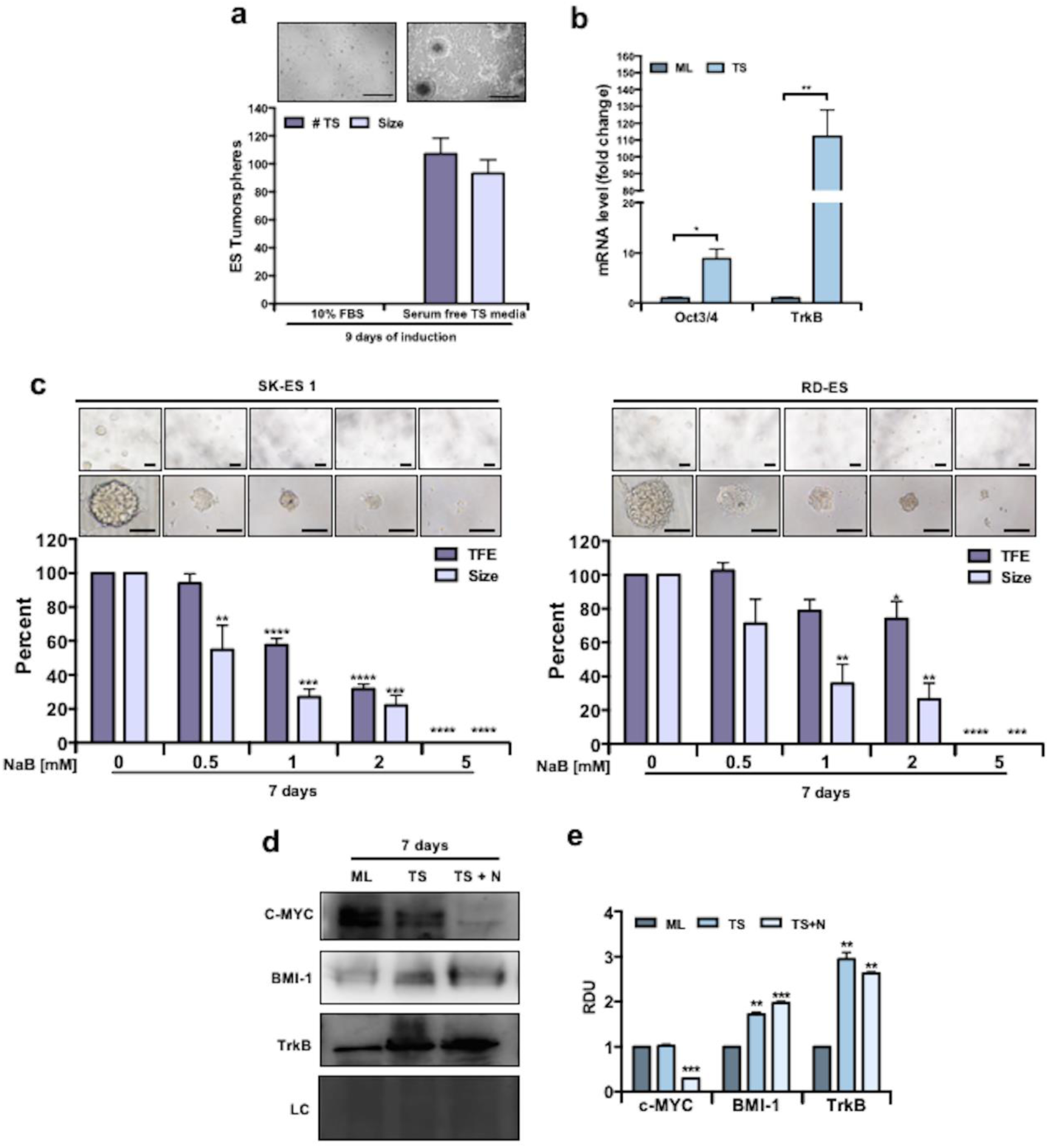
Reduced tumorsphere formation in EWS cells after HDAC inhibition. **a** Percent of number and size of EWS tumorspheres from SK-ES 1 induced with a serum-free tumorspheres medium compared with 10% FBS medium for 9 days; *N* = 3 independent experiments. **b** OCT3/4 and TrkB mRNA levels in SK-ES 1 cells grown in monolayer (10% FBS medium) and tumorspheres (serum-free tumorspheres medium) induced for 9 days. Gene expression levels were normalized by gapdh level and corrected by monolayer; *N* = 3 independent experiments. **c** Representative images from EWS tumorspheres from SK-ES1 (upper left panel) and RD-ES (upper right panel) cells after 7 days of NaB exposure. The percentage of EWS tumorsphere formation efficiency (TFE; i.e., number of tumorspheres normalized by the number of plated single cells and corrected by control – shown in the lower panel - and the size of EWS tumorspheres is shown. Only tumorspheres ≥ 80 µm were analyzed. Scale bars: 200 μm (upper), 100 μm (lower); *N* = 3 independent experiments. **d** Western blot of c-MYC, BMI-1 and TrkB in SK-ES 1 cells in monolayer (ML), Tumospheres (TS) induced for 7 days, and tumospheres exposed to 1 mM of NaB for 7 days (TS + N); *N* = 3 independent experiments. **e** Relative Densitometric Unit (RDU) analyses were normalized by coomassie loading control, and corrected by monolayer. Data in the graphs are shown as mean ± SEM; * *p* < 0.05, ** *p* < 0.01, *** *p* < 0.001, **** *p* < 0.0001 *vs*. controls.

## Discussion

Inhibition of HDAC activity has become an increasingly investigated experimental approach, given its potential to ameliorate the aberrant epigenetic state that underlies a variety of cancers and allow the expression of silenced genes that can promote cell death or differentiation. We found that NaB efficiently inhibits HDAC global activity in EWS cells and decreases EWS-FLI1 mRNA expression, cell growth, long-term proliferation, and colony formation. Similar IC_50_ values for HDAC inhibition and cell growth impairment were observed in both cell lines, leading to G1-phase accumulation. These findings support previous reports of NaB arresting cell cycle, particularly at G1-phase, and inducing cell differentiation and apoptosis in multiple cell lines [29-31].

SK-ES-1 and RD-ES cells are both derived from male patients of similar ages (18 and 19 years, respectively), and both share features including the type of fusion to the exons 7 from EWS and 5 from FLI1 (type 2), p53 missense mutation that result in a nonfunctional p53 protein, lack of p21, and high p16^INK4a^ expression [32, 33]. Overexpression of cyclins D (cyclin D1, D2, and D3) can drive aberrant cell cycle regulation and signaling in many human malignant tumors, including EWS tumors and RD-ES cells [34, 35]. Inhibition of HDAC by NaB resulted in a decrease of cyclin D1 protein levels in both cell lines, similarly to what was previously reported in neuroblastoma [36]. Expression levels of cyclin D1, a G1-phase regulator protein, may represent an important component that controls growth arrest prior to cell differentiation in EWS cells.

Polyploidy induction is a proposed mechanism by which HDis control tumor cell growth. For instance, SAHA induced polyploidy in human colon cancer and breast cancer cells, leading to senescence, particularly in cells harboring defective p21WAF1 or p53 [37]. In contrast, we observed a significant reduction in the proportion of cells showing polyploidy after NaB exposure. HDAC inhibition induced changes in pathways related to cell proliferation and survival in EWS cells, including components of neurotrophin signaling. In other types of pediatric and neuroendocrine cancers, differential neurotrophin and Trk expression is associated with patient prognosis and tumor properties including angiogenesis, metastasis, and chemosensitivity [38]. In EWS, we have recently shown enhanced expression of Trk in tumors, and reduced cell growth and increased chemosensitivity after Trk inhibition [39]. Here we show that NaB exposure in EWS cells resulted in a decrease in NGF and BDNF protein levels, accompanied by reductions in components of the ERK and AKT pathways. Given that neurotrophins are known to mediate neuronal survival via activation of ERK, phosphatidylinositol 3-kinase (PI3K) and phospholipase C*γ* (PLC-*γ*) [40], the findings are consistent with the possibility that inhibition of global HDAC activity suppresses Trk-mediated survival pathways in EWS cells.

We also observed a NaB-induced decrease in the levels of the mitotic marker H3 phos-S10. During mitosis in late G2 and M phase of cell cycle, post-translational modification phosphorylation on serine 10 of histone 3 (H3 phos-S10) is taken as a hallmark of condensed chromatin [41]. High levels of H3 phos-S10 in oncogene-transformed cells and human cancer cells are associated with amplified activation of the Ras-mitogen-activated protein kinase (MAPK)-MSK1 pathway [42]. Moreover, H3 phos-S10 overexpression is associated with chromosome instability that plays a role in the maintenance of ploidy [43] and carcinogenesis [44]. There is a crucial role of phosphorylation on serine 10 of histone 3 at the p21 promoter to activate p21 gene through HDAC inhibition by trichostatin A [45]. Moreover, the application of the mathematical modeling from CPD data described by Silva et al. [26], to evaluate the long-term effects of NaB in the EWS population size, provided a relevant information about the behavior of EWS cells after HDAC inhibition by NaB cells, once we observed a delay in the regrowth of the EWS cell population. In this sense, this mathematical modeling not only contributed to understand the acute effect but also the dynamic of growth kinetic of the surviving population that were not eliminated by the treatment, representing an important tool to study anticancer activities of HDAC inhibitors [19, 20, 26].

Carcinogenesis mediated by cancer stem cells may be considered a sort of epigenetic reprograming where there is loss of expression of specific genes that control differentiation and reactivation of genes specifying stemness [46]. Consistent with this rationale, our results indicated that HDAC inhibition can reduce the expression of several pluripotency-associated genes and this is associated with neurite outgrowth and expression of the neuronal differentiation marker β-III tubulin. Some of the genes which had their expression reduced by NaB in our experiments, including c-MYC and BMI-1, can be upregulated by transcriptional changes induced by EWS/FLI [7, 10, 13, 15, 16, 47-49]. A microarray analysis report revealed that HDAC inhibition by TSA suppresses pluripotency genes including as Nanog, Klf4, Sox2 and Sall4 in embryonic stem cells while activating differentiation-related genes [50]. β-III tubulin is a neuron-specific marker and is expressed in early postmitotic and differentiated neurons [51], hence this differentiation neuronal marker has been used to evaluate neural differentiation in EWS [52, 53]. Our findings provide early evidence consistent with the possibility that HDis can reprogram EWS cells towards neural differentiation, although further experiments are required to characterize the possible neural phenotype of HDi-treated EWS cells.

Sphere formation in cancer cell cultures is a widely used platform for cancer stem cell expansion. Cells from tumorspheres had higher levels of pluripotency-related genes such as OCT3/4 and BMI-1, supporting a stem cell phenotype. In addition, tumorspheres showed higher levels of TrkB compared to cells grown in monolayer, suggesting a possible increase in neurotrophin signaling in EWS stem cells [54]. Importantly, NaB reduced sphere formation, indicating that it is capable of targeting EWS stem cells.

Mechanisms other than histone acetylation may be involved in the effects of NaB in cancer cells. Histone lysine butyrylation is another type of epigenetic post-translational mark that can cooperate or compete with acetylation to promote gene expression programs [55-57]. Increases in butyrylation in neuroblastoma cells after exposure to SAHA have been reported [58]. It should also be noted that NaB has been shown to enhance cAMP levels and stimulate the activation of multiple protein kinase pathways through mechanisms at least partially mediated by effects independent of epigenetic influences, through direct interactions with cAMP/PKA signaling in the cytoplasm [59-61]. Thus, we cannot exclude the possibility that NaB-induced effects on cell machinery components other than HDACs are involved in our findings.

This study has several limitations that will need to be addressed by further experiments. For instance, the mechanistic relationship among the actions of NaB, the reduction in EWS-FLI1 expression, and the reduced expression of the other target genes remains highly speculative. In addition, histone acetylation changes after HDi treatment should be measured. Furthermore, the effects of different HDis should be compared, and confirmed in *in vivo* models.

A schematic model of the biological actions of NaB in EWS is depicted in Fig. 8. This model proposes a mechanism by which NaB acts through HDAC inhibition on transcriptional regulators to change the differentiation status of EWS cells. Together, our results indicate that HDis can reduce EWS growth and survival and lead to a more differentiated state. It is noteworthy that these data are consistent with our previous findings indicating that HDis can impair cell growth and modulate differentiation of cells from other types of childhood pediatric tumors possibly originating from neural stem cells, medulloblastoma and neuroblastoma [62, 63].

**Fig. 8.**
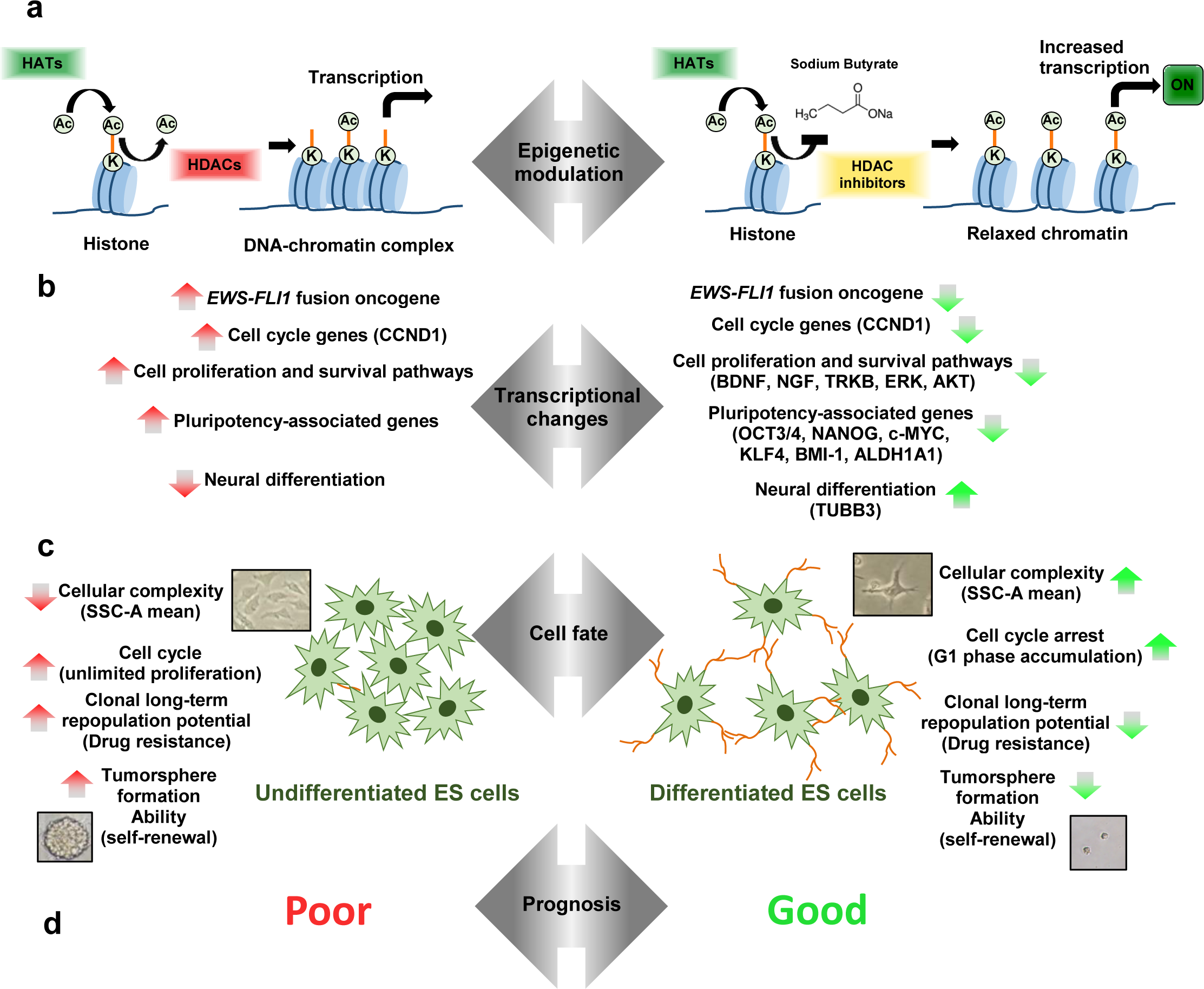
Schematic model depicting biological actions HDAC inhibition by NaB in EWS. **a** Unbalance between the activities of histone acetyltransferases (HATs) and histone deacetylases (HDACs) results in an aberrant epigenetic state that occurs often in EWS cells. HDAC inhibitors such as NaB have been employed to revert this setting by increasing histone acetylation and gene transcription. **b** As a result of epigenetic dysregulation, EWS cells present an aberrant expression of genes associated cell cycle, proliferation, survival, and pluripotency pathways. HDAC inhibition by NaB leads to transcriptional change by suppressing cell cycle, proliferation, survival, and pluripotency-associated genes, whereas de-repressing expression of the neural differentiation protein TUBB3. **c** A poor differentiated phenotype is a feature of EWS. HDAC inhibition change cell fate towards increased cellular complexity, arrested cell growth by G1-phase accumulation, decreased long-term clonal repopulation and reduced tumorsphere formation. **d** Inhibiting HDAC might epigenetically reprogram EWS cells from an undifferentiated to a more differentiated state associated with a better prognosis.

## Acknowledgements

This research was supported by PRONON/Ministry of Health, Brazil (number 25000.162.034/2014-21); the Children’s Cancer Institute (ICI); the National Council for Scientific and Technological Development (CNPq; grant number 303276/2013-4 to R.R.); the Rio Grande do Sul State Research Foundation (FAPERGS; grant number 17/2551-0001 to R.R.); the Coordination for the Improvement of Higher Education Personnel (CAPES; to B.K.S.); the Clinical Hospital institutional research fund (FIPE/HCPA); and the Center for Cancer Research, National Cancer Institute, National Institutes of Health (C.J.T.).

